# Genomic profiling of PML bodies reveals transcriptional regulation by PML bodies through the DNMT3A exclusion

**DOI:** 10.1101/729483

**Authors:** Misuzu Kurihara, Kagayaki Kato, Chiaki Sanbo, Shuji Shigenobu, Yasuyuki Ohkawa, Takeshi Fuchigami, Yusuke Miyanari

## Abstract

The promyelocytic leukaemia (PML) body is a phase-separated nuclear structure involved in various biological processes, including senescence, and tumour suppression^1^. PML bodies consist of various proteins, including PML proteins and several chromatin regulators^2,3^ and physically associate with chromatin^4,5^, implying their crucial roles in particular genome functions. However, their roles in transcriptional regulation are largely unknown. Here, we developed APEX-mediated chromatin labelling and purification (ALaP), to identify the genomic regions associated with PML bodies. We find that PML bodies associate with active regulatory regions across the genome and prominently with a ∼300 kb of the short arm of the Y chromosome (YS300) in mouse embryonic stem cells (mESCs). The association with YS300 is essential for the transcriptional activities of neighbouring Y-linked cluster genes. Mechanistically, we show that PML bodies play a novel role in 3D nuclear organization by providing specific nuclear spaces that the de novo DNA methyltransferase DNMT3A cannot access, which results in the robust maintenance of the hypo-methylated states at the Y-linked gene promoters. Our study underscores a new mechanism for gene regulation in the 3D-nuclear space and provides insights into the functional properties of nuclear structures for genome functions.

## Main

Previous studies indicated that PML bodies tend to localize close to actively transcribing gene loci^6,7^. Several other works reported that the association between PML bodies and specific gene loci could affect the gene expression activities^8–11^. These evidences prompted us to profile the genomic regions associated with PML bodies and investigate the regulatory potential of PML bodies on chromatin functions. Chromatin immunoprecipitation sequencing (ChIP-Seq) is a powerful approach for profiling protein binding to chromatin. However, PML bodies are insoluble, and therefore ChIP-seq analyses are not adapted^12,13^. APEX2 is an engineered peroxidase that displays superior activities in living cells^14^, where it catalyses the oxidization of biotin-phenol to generate a radical form in an H_2_O_2_-dependent manner and results in biotinylation of proteins in tight spaces (< 20 nm in radius). APEX2 has been used for the identification of proteins proximal to target proteins or RNA by being combined with mass spectrometry^15–18^. Compared with other proximity labelling techniques for chromatin studies such as bioChIP-seq^19^, imuno-Trap^20^ and TSA-seq^21^, the quick reaction (1 min) by APEX2 in living cells is suitable to address the dynamic association between nuclear domains and chromatin.

To assess whether ALaP enables us to enrich the genomic sequences, we applied it to analyze chromatin binding of NANOG, a transcription factor well characterized in mESCs^22,23^, as a proof of principle. We established mESCs in which APEX2 is knocked in the *Nanog* allele as a C-terminal fusion protein (Nanog-APEX)(Supplementary Fig. 1a-c). NANOG-binding sites identified previously^24^ were efficiently enriched in the biotinylated fraction (Supplementary Fig. 1d), indicating that ALaP can be used to address the interaction between a protein of interest and its chromatin targets. We next applied ALaP to identify the genomic loci associated with PML bodies using mESCs in which APEX2 is knocked in the *Pml* locus as an N-terminal fusion protein (APEX-Pml) (Supplementary Fig. 2a). As expected, APEX2 proteins were specifically recruited to PML bodies (Supplementary Fig. 2b). Optimizations of the ALaP workflow resulted in specific labelling of spaces proximal to PML bodies (Fig. 1a,b, Supplementary Fig. 2c-e). In agreement with the known localization of telomeres to PML bodies in mESCs (Fig. 1c)^3^, the telomeric sequence was significantly enriched by ALaP (Fig. 1d). In contrast, we did not find the enrichment by ChIP with anti-PML antibody in wild-type mESCs and with anti-GFP or anti-HA antibodies in GFP-HA-PML-knockin mESCs (Fig. 1d, and Supplementary Fig. 2f). These results indicate that ALaP is useful for identifying genomic loci proximal to PML bodies, which cannot be addressed by conventional ChIP.

**Fig 1.**
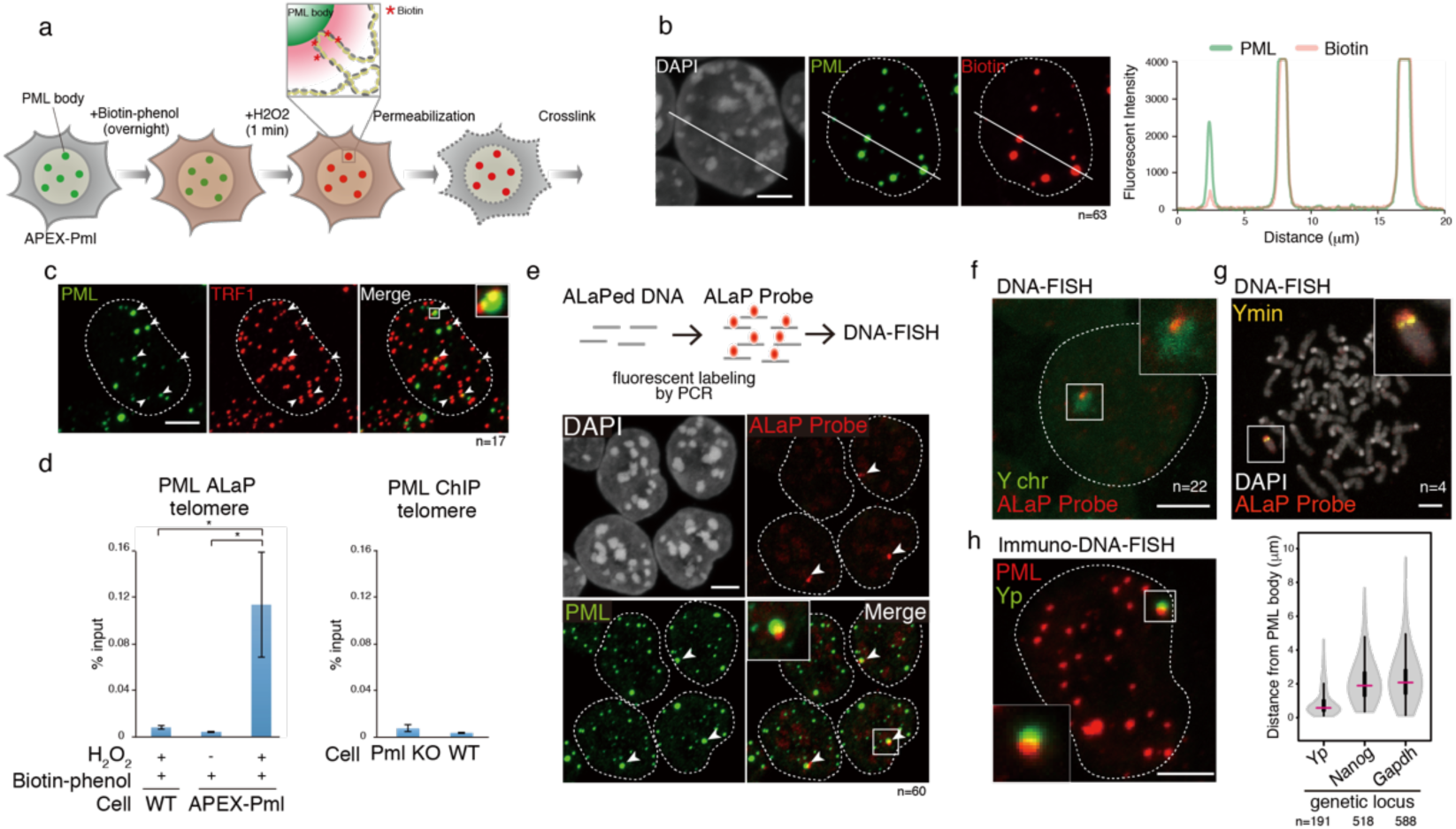
Development of ALaP. (a) Schematics of workflow for labeling of PML bodies by APEX reaction. (b) Representative images of biotinylated fractions (red), PML proteins (green) and DAPI (gray) after APEX reaction in APEX-Pml mESCs. Intensities of PML and biotin along a white line is shown in a right panel. Dashed lines indicate periphery of nucleus. Scale bars, 5 µm. n, number of cells analysed. (c) Representative images of immunostaining for PML (green) and TRF1, a telomere marker (red). PML bodies adjacent to telomeres are indicated with arrowheads. (d) qPCR analysis for enrichment of telomeric sequence in ALaP (left) and conventional ChIP experiment with anti-PML antibody (right). ALaP were conducted with or without H_2_O_2_ treatment. Experiments were performed with indicated cell lines. The mean ± s.d. of three independent biological replicates is shown. Statistical comparisons were performed by Dunnett test. *, *P*<0.05. (e) Workflow for preparation of ALaP probe for DNA-FISH (top). Representative Immuno-DNA-FISH images for ALaP probe (red), PML (green) and DAPI (gray) are shown in bottom panels. Arrowheads indicate bright spots of DNA-FISH. (f) A representative image of DNA-FISH with ALaP probe (red) and painting probe for Y chromosome (Y chr, green) in mESCs. (g) DNA-FISH for centromere of Y chromosome (Ymin, yellow) and ALaP probe (red) with DAPI stained (gray) on chromosome spreads of mESCs. (h) A representative image of Immuno-DNA-FISH with BAC probe for Yp (green) and PML (red)(left). Distribution of distance between indicated genetic locus and the closest PML body is shown in a violin plot with median (red line), box (25th and 75th percentiles), and whiskers (1.5 times the interquartile range from the 25th and 75th percentiles).

**Fig 2.**
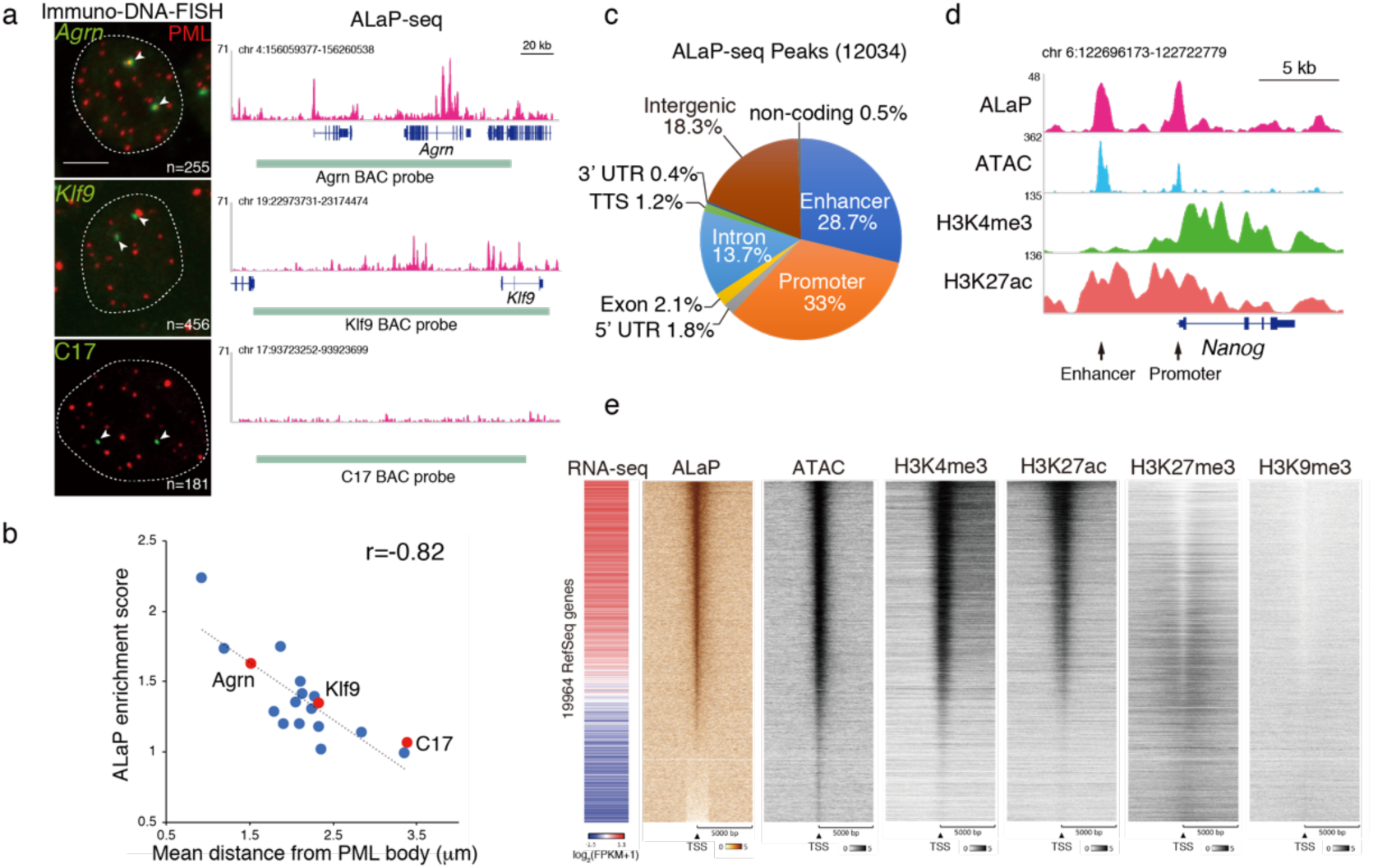
ALaP-seq uncovered association of PML bodies with regulatory regions of active genes. (a) Representative images of Immuno-DNA-FISH for BAC probes of indicated genetic loci (green) and PML proteins are shown in a left panel. Dashed lines indicate periphery of nucleus. Arrowheads indicate spots of DNA-FISH. Scale bars, 5 µm. n, number of cells analysed. ALaP-seq tracks along corresponding genomic loci are shown in right panels with green lines showing position of indicated BAC probes. (b) Correlation between enrichment of ALaP-seq reads and mean distance between genomic locus and the closest PML body analyzed by Immuno-DNA-FISH. The Pearson correlation coefficient (r) is indicated on the graph. Genetic loci represented in (a) were colored in red. (c) A pie chart of genome-wide distribution of ALaP-seq peaks on indicated genomic elements including predicted enhancer and promoter (±1 kb to TSS). (d) Genome browser track displaying enrichment of ALaP-, ATAC-, and ChIP-seq for H3K4me3 and H3K27ac along *Nanog* locus. *Nanog* promoter region and enhancer are indicated with arrowheads, respectively. (e) Heat maps of ALaP-seq (orange) and indicated epigenetic profiling (black) aligned at ± 5000 bp of TSS of 19964 individual RefSeq genes, sorted by ALaP-seq levels. Expression levels of corresponding genes analyzed by RNA-seq is shown in a left panel.

To further evaluate our approach, we used DNA enriched by ALaP as a fluorescence probe (ALaP probe) for DNA-FISH. Bright fluorescent spots for the ALaP probe were detected adjacent to PML bodies (Fig. 1e), which was not observed with a negative control probe prepared under H_2_O_2_ omitted conditions (data not shown), strongly suggesting that genomic loci proximal to PML bodies are specifically enriched by ALaP. Remarkably, the number of the ALaP spots was single per nucleus (Fig. 1e), indicating that genomic regions interacting with PML bodies may be on the X or Y chromosome (chr), as mESCs used are male diploid. Further DNA-FISH analyses revealed a specific localization of the short arm of Y chr (Yp) onto PML bodies (Fig. 1f,g). Additionally, the 3D distance between the Yp and PML body is significantly shorter than other genetic loci (Fig. 1h), demonstrating that the Yp locus frequently localized in the vicinity of PML bodies in mESCs.

We performed deep sequencing analysis of DNA enriched by the PML-ALaP to profile the genomic loci proximal to PML bodies. In agreement with our qPCR results (Fig. 1d), the telomeric sequence was significantly enriched in an APEX reaction-dependent manner (Supplementary Fig. 3a). We asked whether our ALaP-seq recapitulates the association between genomic loci and PML bodies by conducting 3D Immuno-DNA-FISH analyses with 18 BAC probes encoding different genomic regions displaying the diverse ALaP enrichments. Remarkably, the enrichment of ALaP-seq for each BAC probe is highly correlated with mean distances between PML bodies and the corresponding genomic loci (Fig. 2a,b). Thus, we conclude that ALaP-seq allows us to explore genomic loci proximal to PML bodies.

**Fig 3.**
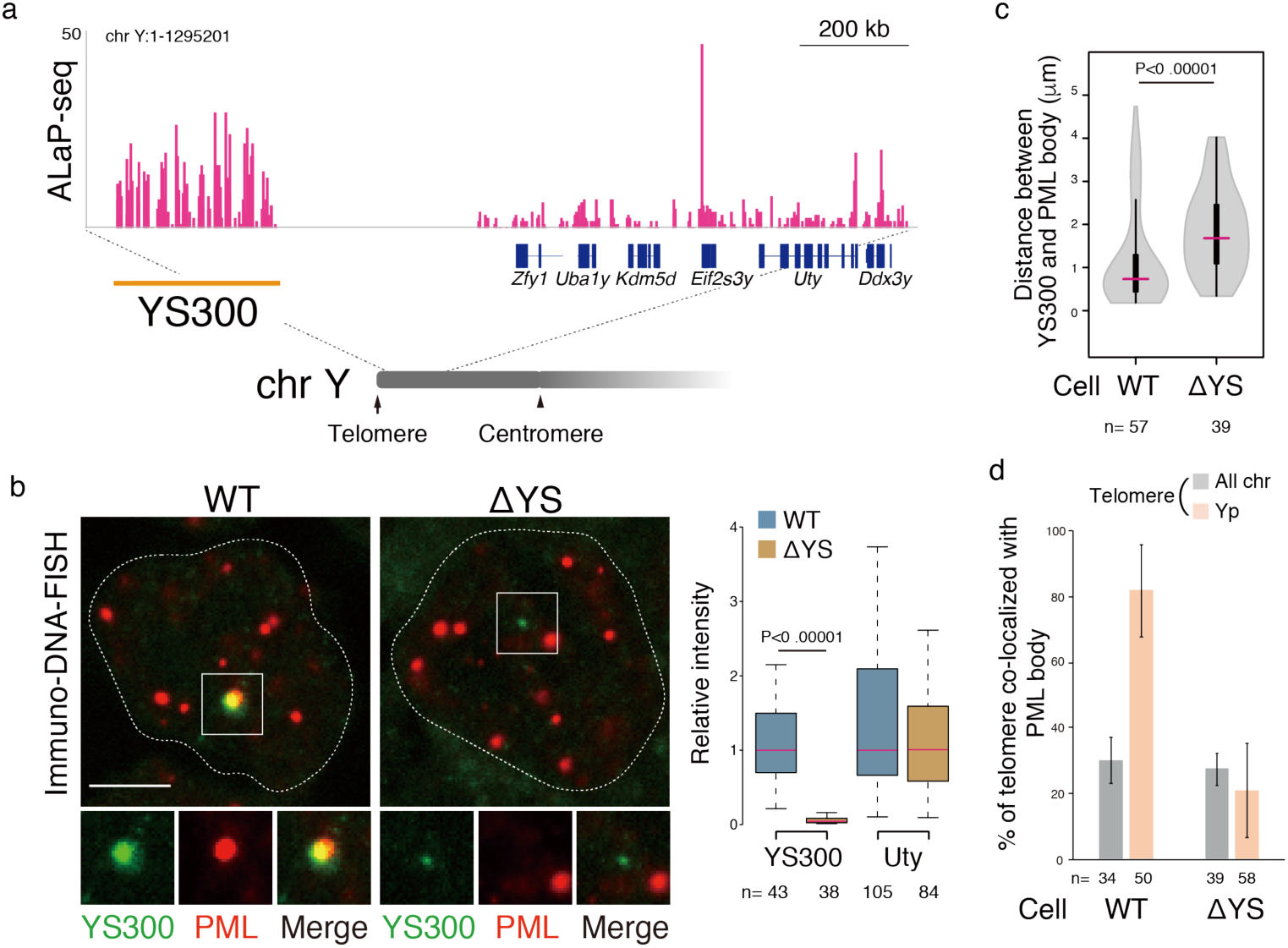
YM300 is a prominent region frequently associated with PML bodies. (a) Enrichment of ALaP-seq along indicated genomic region of Yp. (b)Representative images of immune-DNA-FISH for PML proteins (red) and YS300 (green) in WT and ΔYS mESCs are shown in left panels. Close-up views of boxed area are shown at bottom. Dashed lines indicate periphery of nucleus. Scale bars, 5 µm. Box plots show distribution of relative intensity of YS300 and *Uty* locus in WT and ΔYS mESCs (right). n means number of cells analyzed. (c) Distribution of distance between YS300 and the closest PML body in WT and ΔYS mESCs is shown in a volcano plot. (d) Percentage of telomere of Yp or all chromosomes co-localized with PML bodies in WT and ΔYS mESCs is shown. Data for Yp telomere was calculated based on immuno-DNA-FISH with PML and BAC probe for YS300, and one representing all telomeres are analyzed by immunostaining for TRF1 and PML. Statistical comparisons were performed by Mann-Whitney U test.

ALaP peaks were notably enriched at regulatory regions, including core promoters and enhancers across the genome, as represented by the *Nanog* locus (Fig. 2c,d, and Supplementary Fig. 3b,c). The enrichment of ALaP-seq within TSS proximal regions are highly correlated with the expression levels of the associated genes and the epigenetic signatures of active promoters, including accessible chromatin (as determined by ATAC-seq), H3K27ac and H3K4me3. In contrast, repressive marks, including H3K9me3 and H3K27me3 were depleted (Fig. 2e, and Supplementary Fig. 3d-e). Taken together, these results strongly suggest that PML bodies associate primarily with regulatory regions of active genes in a genome-wide manner. This is consistent with previous studies showing that actively transcribing loci were detected in the vicinity of PML bodies^4,6^. Unlike the broad chromatin peaks observed with other nuclear domains, such as nuclear lamina and speckles^21,25^, ALaP-seq displayed sharp peaks (Supplementary Fig. 3f), suggesting that PML bodies associates within a narrow band of chromatin.

While post-translational modification by SUMO is involved in recruiting a target protein to PML bodies^26,27^, comparing ALaP-seq and publicly available ChIP-seq datasets for SUMO-1 and SUMO-2/3^28^ did not detect a significant correlation (Supplementary Fig. 4), indicating that the association of PML bodies with chromatin is likely to be independent of the global pattern of SUMOylation on chromatin.

**Fig 4.**
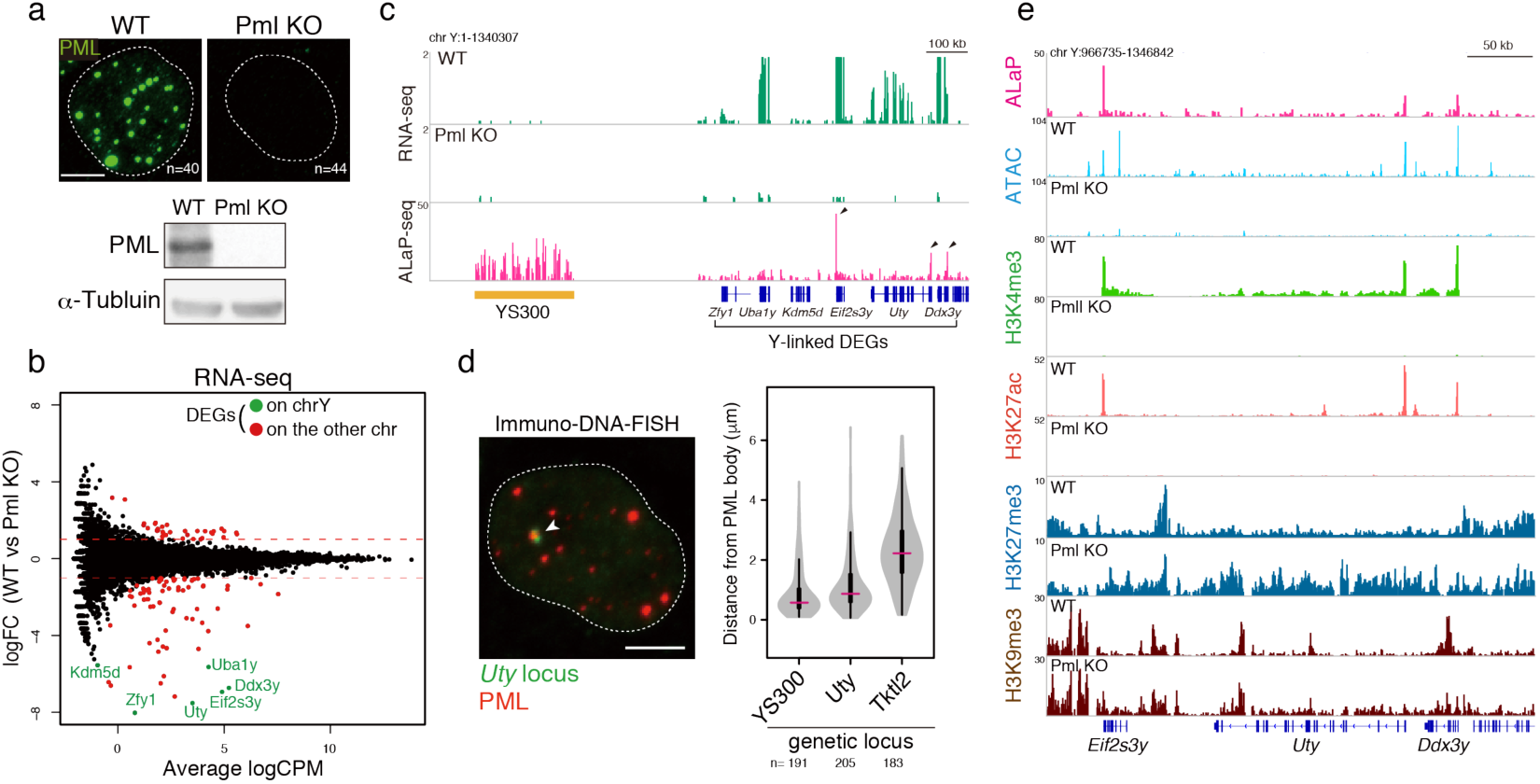
PML bodies positively regulate expression of Y-linked cluster genes. (a) Immunostaining images of PML (green) in indicated cell lines are shown (top). Dashed lines indicate periphery of nucleus. Scale bars, 5 µm. n, number of cells analysed. Western blots of PML protein and α-Tubulin with corresponding cells are shown in bottom panels. (b) A MA plot of RNA-seq representing Pml KO versus WT mESCs. DEGs with a FDR < 0.001 and |log_2_FC|>1 are highlighted in colors. Y-linked DEGs are labeled in green. (c) RNA-seq and ALaP-seq tarack in WT and Pml KO mESCs for indicated Yp loci. Arrowheads indicate peaks on promoter regions of corresponding genes. (d) A representative Immuno-DNA-FISH image for PML (red) and *Uty* locus (green) (left). An arrowhead indicates spot of DNA-FISH. Distribution of distance between indicated genetic locus and the closest PML body is shown in a volcano plot. (e) Genome browser tracks showing enrichment of ALaP-, ATAC-, and ChIP-seq for H3K4me3, H3K27ac, H3K27me3, and H3K9me3 along Y-linked clustered gene loci in WT and Pml KO mESCs.

We next addressed whether the ALaP-seq recapitulated our DNA-FISH result in which Yp was specifically stained with the ALaP probe (Fig. 1e). Since the mouse Y chromosome encodes a variety of Y-specific repetitive sequences^29^, we analyzed ALaP-seq for association with repetitive elements. Interestingly, around the 300 kb block of the sub-telomeric region in Yp (hereafter referred to as YS300) displayed the significant enrichments (Fig. 3a). YS300, indeed, consists of specific inverted repeats that are not homologous to other sequences in the genome and are flanked by unassigned sequences in the 5’ and 3’ regions (Supplementary Fig. 5a). YS300 displays significantly higher ALaP-seq enrichment than other 300 kb genomic bins (Supplementary Fig. 5b). Other genomic regions also display higher ALaP enrichments, in contrast to our DNA-FISH data with the ALaP probe, in which only a single spot was detected in each nucleus. This discrepancy might be due to technical differences between the DNA-FISH and deep sequencing. Furthermore, DNA-FISH spots for the ALaP probe completely overlapped with those of the BAC probe that encompasses the YS300 sequences but not with those of the other probes tiled at Yp (Supplementary Fig. 5c), suggesting that YS300 is the genomic loci associated with PML bodies. Indeed, among all 18 genomic loci analysed by DNA-FISH, YS300 exhibits the shortest distance to PML bodies. Colocalization analysis between PML bodies and telomeres revealed that the telomeric side of the Yp is associated with a PML body in 80% of mESCs, which is much more frequent than the 30% association of other telomeres (Fig. 3d). Furthermore, the Yp locus preferentially associates with a large PML body (Supplementary Fig. 5d). Taken together, YS300 is a hotspot associated with PML bodies in mESCs. To address the dependency of YS300 to the PML body-association, we established a mutant mESCs in which the majority but not all of the YS300 sequence was deleted by CRISPR/Cas9 (ΔYS) (Fig. 3b, and Supplementary Fig. 5a). Remarkably, the deletion resulted in the dissociation of the Yp locus from PML bodies (Fig. 3b,c). In addition, the frequency of the colocalization between the telomeric side of the Yp and PML bodies was drastically reduced to the comparable level of other telomeres (Fig. 3d), suggesting that YS300 is crucial for anchoring or nucleation of PML bodies on the Yp locus.

**Fig 5.**
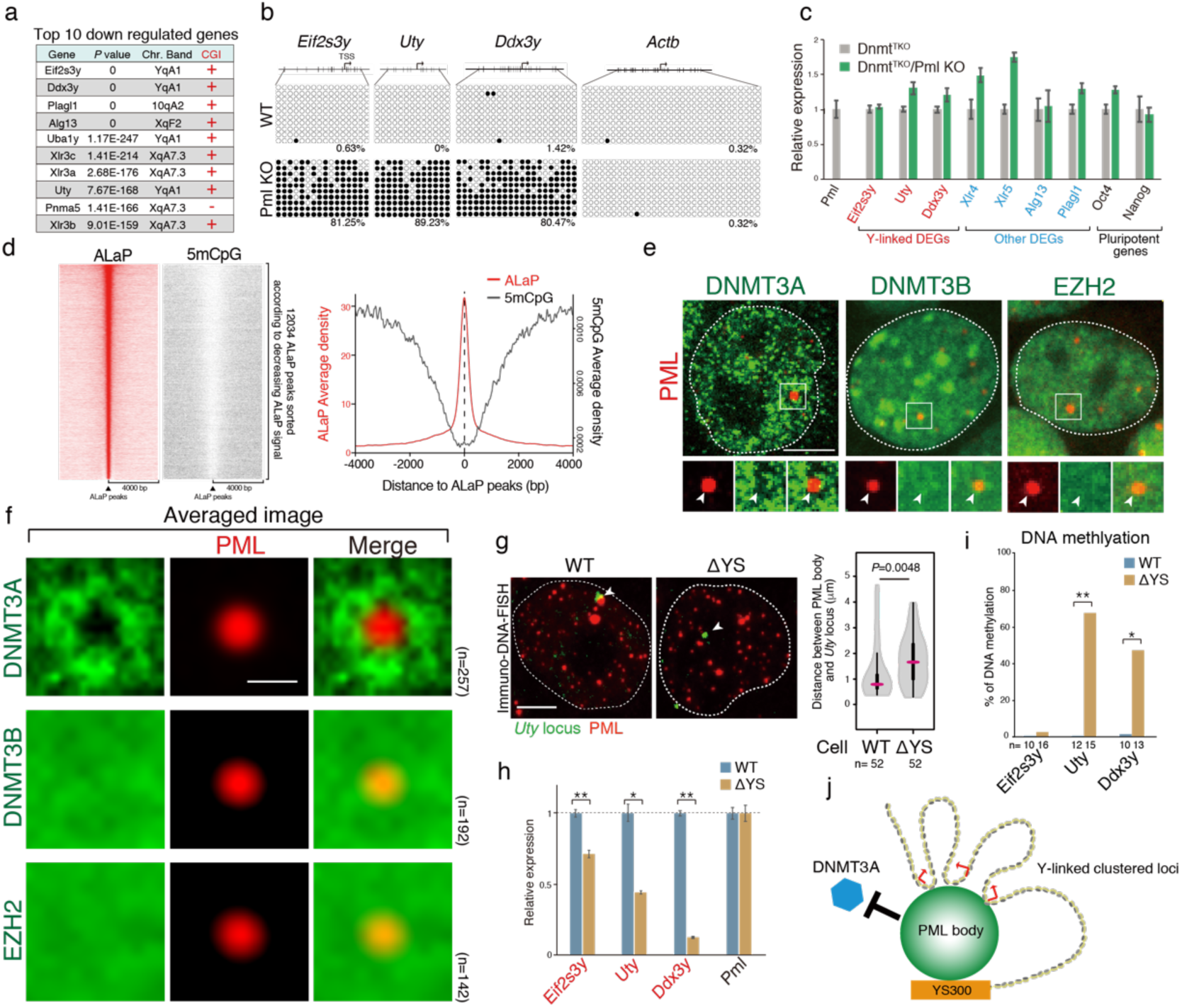
PML bodies exclude Dnmt3a to maintain the transcriptional activity of the Y-linked clustered genes. (a)Top 10 downregulated DGEs is shown with *P*-value, chromosomal band position, CpG islands (CGI) at their promoters (within ±1 kb to TSS). (b) DNA methylation status of promoter regions of indicated genes analyzed by bisulfite sequencing. Position of a CpG dinucleotide is represented by vertical lines. Open circle, unmethylated CpG; closed circle, methylated CpG. The percentages of methylated CpG are indicated at bottom of corresponding panels. (c) Relative expression of indicated genes in Dnmt^TKO^ and Dnmt^TKO^ lacking PML (Dnmt^TKO^/Pml KO) mESCs. qRT–PCR data are presented relative to Dnmt^TKO^ cells. The mean ±s.d. of three independent biological replicates is shown. (d)Heat map of ALaP-seq (red) and DNA methylation (5mCpG) profiling (black) aligned at ± 4000 bp at centers of ALaP-seq peaks, sorted by ALaP-seq signals (left). An aggregation plot showing enrichment of 5mCpG and ALaP-seq around ALaP peaks (right). (e) Representative immuno fluorescence images for indicated proteins in mESCs. Dashed lines indicate periphery of nucleus. Insets on bottom show enlarged images of the boxed regions. Arrowheads indicate position of PML bodies. (f) Averaged image projection of immunostaining for indicated proteins (green) centered at PML bodies (red) are shown. n, number of PML bodies analyzed for calculation of the averaged signals. (g) Representative Immuno-DNA-FISH images for PML proteins (red) and *Uty* locus (green) in WT and ΔYS mESCs (left). Arrowheads indicate spots of DNA-FISH. Distribution of distance between *Uty* locus and the closest PML body in WT and ΔYS mESCs is shown in a volcano plot (right). Statistical comparison was performed by Mann-Whitney U test. (h) Relative expression of indicated genes in WT and ΔYS mESCs was analyzed by qRT-PCR and normalized to WT. The mean ±s.d. of three independent biological replicates is shown. Statistical comparisons were performed by t-test. *, *P*<0.01; **, *P*<0.001. (i) DNA methylation levels at promoters of indicated Y-linked DEGs in WT and ΔYS mESCs. Statistical comparisons were performed by Mann-Whitney U test. *, *P*<0.01; **, *P*<0.001. (j) A model for regulation of the Y-linked cluster genes by PML bodies. Scale bars, 5 µm in (e, g), 1 µm in (f)

We asked whether the PML body-association could contribute to transcriptional gene regulation by performing RNA-seq analysis with wild-type and PML-knockout mESCs (Pml KOs) which lacked PML bodies (Fig. 4a). While the PML deletion did not result in a general defect of transcription of PML-associated loci (4971 genes) (Supplementary Fig. 6a), we identified 104 differentially expressed genes (DEGs, |log_2_FC| >1, FDR < 0.001) in Pml KO (Fig. 4b). Among these DEGs, 70 genes were downregulated to a greater extent than the upregulated genes, indicating that PML mainly display positive effects for gene expression. A gene ontology analysis revealed that the DEGs related to spermatid development and telomere maintenance were enriched (Supplementary Fig. 6b).

Interestingly, the depletion of PML resulted in a marked reduction in the expression levels of six Y-linked genes (Fig. 4c), without affecting their paralogues in the X chromosome or of pluripotent genes (Supplementary Fig. 6c). We confirmed that the down regulation of Y-linked genes and other DEGs was rescued by exogenous PML gene expression, while the effect was limited (Supplementary Fig. 6d). To our surprise, all of these Y-linked DEGs are indeed clustered together at a 600 kb distance from YS300 (Fig. 4c). Furthermore, enrichment of ALaP-seq was observed on promoters of these cluster genes (Fig.4c), which was consistent with the adjacent localization of PML bodies to the *Uty* locus, as shown by DNA-FISH (Fig. 4d). In addition, nascent RNA transcripts of these Y-linked genes were predominantly transcribed in the vicinity of PML bodies (Supplementary Fig. 6e). Taken together, these results suggest a model whereby the association between PML bodies and the Y-linked cluster genes regulate their expression in mESCs.

To address the mechanisms whereby PML bodies regulate Y-chromosome-linked gene expression, we first asked whether misregulation of genes is due to their subtelomeric location rather than the specific association with PML bodies. However, analysis of the genomic location of all DEGs revealed that they were spread across chromosome arms, and were not enriched on subtelomeric regions (Supplementary Fig. 7a). Only 12 out of all 329 subtelomeric genes (3.6%) were affected upon PML depletion. The impact on 6 Y-linked DEGs was outstanding compared with the other 6 subteromeric DEGs (Supplementary Fig. 7b,c). These results collectively suggest that the Y-linked cluster locus is a prominent site specifically regulated by PML bodies. Since *Eif2s3y, Uty*, and *Ddx3y* are highly expressed among 6 Y-linked genes in mESCs, and show broad expression in other tissues^30^, we selected them for subsequent analyses.

We next investigated the influence of PML bodies onto the chromatin signatures of Y-linked cluster gene loci. ChIP- and ATAC-seq analyses showed that loss of PML bodies induced a drastic reduction in transcriptionally permissive euchromatic signatures, including those of H3K4me3 and H3K27ac and chromatin accessibility, at their promoters, and instead, the repressive epigenetic mark H3K27me3, but not H3K9me3, was increased (Fig. 4e). These findings indicate that the PML body-mediated transcriptional modulation of the Y-linked cluster genes coincides with alterations in epigenetic signatures.

To understand how PML bodies regulates the expression level of these cluster genes, we first focused on H3K27me3, because their entire gene loci were covered with H3K27me3 upon PML depletion (Fig. 4e). mESCs lacking Ezh2 (Ezh2 KOs), a major enzyme responsible for H3K27 trimethylation, displayed a global reduction of H3K27me3, as reported previously^31^ and maintained the PML body-association with YS300 (Supplementary Fig. 8a,b). We found that loss of PML bodies in Ezh2 KOs led to a marked reduction of Y-linked genes (Supplementary Fig. 8c), as we observed in PML KOs (Fig. 4), which demonstrated that the repression of Y-linked genes upon the PML depletion depends on other factors rather than H3K27me3.

Interestingly, 9 out of the top 10 downregulated DEGs contain CpG islands (CGIs) in their promoters (Fig. 5a). Therefore, we next assessed DNA methylation levels at the promoter in *Eif2f3y, Uty* and *Ddx3y*, and found that their hypo-methylated states in wild-type mESCs were drastically converted to hyper-methylated states in Pml KOs (Fig. 5b). In addition, promoter regions in other DEGs, such as *Xlr4, Xlr5*, and *Plagl1*, also had higher levels of DNA methylation in Pml KOs (Supplementary Fig. 8d). These results suggest that DNA methylation might be crucial to regulate the Y-linked cluster genes and other DEGs. To address this possibility, we analysed DNA methyltransferase triple knockout cells (Dnmt^TKO^)^32^. We verified that the expression levels of the Y-linked genes and the PML body-association with YS300 were comparable to those in wild-type mESCs (Supplementary Fig. 8b,e). Strikingly, depletion of PML in the Dnmt^TKO^ cells did not lead to the repression of the Y-linked genes and other DEGs (Fig. 5c), which was in sharp contrast to that observed in PML KO and Ezh2/PML-double knockout mESCs (Fig. 4 and Supplementary Fig. 8c). These results strongly indicate that DNA methylation is involved in the PML body-mediated transcriptional regulation of the Y-linked genes. In the other words, the association with PML bodies could protect the Y-linked genes from DNA methylation. In line with this, a comparison between ALaP-seq and available whole genome bisulfite sequencing data revealed that ALaP-peaks exhibited low DNA methylation states compared to the levels in surrounding regions (Fig. 5d). In addition, reads enriched with ALaP reads on CpG islands were negatively correlated with their DNA methylation levels (Supplementary Fig. 9a-c), suggesting that PML bodies are preferentially associated with hypo-methylated regions.

We next performed immunostaining for DNMT3A and DNMT3B, de novo DNA methyltransferases (Supplementary Fig. 9d). DNMT3A displayed heterogeneous speckle localization throughout the nucleoplasm (Fig. 5e), consistent with previous reports^33,34^. DNMT3B accumulated in DAPI dense heterochromatin, and diffusively localized throughout the nucleoplasm. Strikingly, we observed that DNMT3A was consistently excluded from PML bodies (Fig. 5e). Averaged image projections centered on PML bodies clearly revealed that the intensity of DNMT3A was lower at the centre of the PML bodies than in the surrounding area, contrary to the homogeneous distribution of DNMT3B and EZH2 (Fig. 5f). This suggests that PML bodies provides specific nuclear spaces that DNMT3A cannot access.

To investigate how the PML body-association affects to the DNA methylation status, we analysed ΔYS mESCs and found that deleting YS300 led to dissociation of the *Uty* locus from PML bodies (Fig. 5g). In addition, YS300 and the *Uty* locus almost always share the same PML body in wild-type mESCs (Supplementary Fig. 9e). These results strongly suggest that anchoring of a PML body by YS300 promotes the association of Y-linked gene loci with PML bodies. Importantly, we observed significant downregulation of Y-linked cluster genes in ΔYS, and its effect was correlated with their genetic distance from telomeres (Fig. 5h). In line with this, the repression was accompanied by the significant acquisition of DNA methylation at their promoters, especially in *Uty* and *Ddx3y* (Fig. 5i), indicating that the association between a PML body and the Y-linked loci is crucial for their transcriptional activity through suppression of DNA methylation. These results collectively provide further support for PML bodies playing a role in maintaining a hypo-methylated state to regulate the transcriptional activity of Y-linked genes by preventing DNMT3A access (Fig. 5j).

In this study, we propose that PML bodies play a crucial role in 3D nuclear organization to regulate the transcriptional activity through exclusion of DNMT3A but not DNMT3B. This finding aligns with selective features of PML bodies in which some specific proteins are accumulated and others are excluded or evenly distributed^35^. While SUMO-dependent accumulation of proteins to PML bodies has been extensively studied^27,36^, the rules by which molecules are excluded from PML bodies are largely unknown. DNMT3A and DNMT3B have different interacting partners^37^ and display unique binding specificities to the genome. While DNMT3B selectively targets gene bodies, DNMT3A2, a DNMT3A isoform predominantly expressed in mESCs, has no binding preference^38^. These specific features could contribute to the selective access to PML bodies.

A key question concerns the driving force that recruits PML bodies to specific genomic regions. Sequence motif analysis of PML-ALaP peaks did not reveal any particular enrichments (data not shown). One of the highly possible factors involved in this recruitment is SUMO modification. SUMOylation of the telomere-binding proteins TRF1 and TRF2 is required for the recruitment of telomeres to PML bodies^38,39^; however, we did not observe a positive correlation between the SUMO-associated and PML body-associated chromatin in mESCs. Considering preferential association of PML bodies with active regulatory regions, transcription activity could be a key for the recruitment, as proposed previously^40^.

We show that the Y-linked cluster genes, including *Eif2s3y, Uty*, and *Ddx3y* are positively regulated by the association of PML bodies. Several reports have suggested that all these genes are implicated to play roles in sexual dimorphism^30,41–43^. Indeed, the downregulation of these genes upon the PML-depletion was also observed in mouse embryonic fibroblasts (MEFs) (Supplementary Fig. 6f)^13^, suggesting that these genes might be robustly regulated by PML bodies. It will be interesting to investigate whether PML bodies could contribute to the sex differences, which has never been addressed yet.

## Materials and Methods

### Cell culture

All the mouse ES cell lines were cultured in DMEM (Wako, 043-30085) supplemented with 20% FBS, GlutaMAX (Gibco), MEM Non-Essential Amino Acids (Gibco), Leukemia inhibitory factor, 1 mM sodium pyruvate, penicillin/streptomycin, 0.1 mM 2-mercaptoethanol, 3 mM CHIR99021 and 1 mM PD0325901. Cells were cultured at 37 °C under 5% CO_2_ in air.

### Preparation of cell lines

For generation of APEX-Pml-knockin cells, wild-type ES (E14) cells were electroporated with a targeting vector composed of APEX2 and a puromycin resistance gene as shown in Supplementary Fig. 2a. For preparation of Nanog-APEX-knockin cells, wild-type mESCs were transfected with a targeting vector (illustrated in Supplementary Fig. 1a), and pX300A-G+Nanog expressing Cas9 and sgRNAs targeting the Nanog allele by using Lipofectamine 2000 (Invitrogen, 11668019). For preparation of EGFP-HA-Pml-knockin cells, mESCs were transfected with a targeting vector pGb_PMLlxP_EGFP, and pYMCR2_Gb_PMLTSS expressing Cas9 and sgRNAs targeting the vector and PML allele by using Lipofectamine 2000. Cells were selected with 2 µg/ml of puromycin for a week. The targeted clones were screened by genotyping PCR, western blot, and southern blotting (data not shown).

For establishment of Ezh2 knockout mESCs, E14 was transfected with pSpCas9(BB)-2A-Puro (PX459) V2.0 (Addgene, 62988) harboring sgRNAs targeting Ezh2 allele by using Lipofectamine 2000 and selected with 3 µg/ml of puromycin for 2 days. The resultant single colonies were screened by immunofluorescence staining with anti-EZH2 antibody (data not shown).

For establishment of PML knockout cells, E14 was transfected with pSpCas9(BB)-2A-Puro (PX459) V2.0 or pECas9B harboring sgRNAs targeting *Pml* allele by using Lipofectamine 2000 and selected with 3 µg/ml of puromycin or 20 µg/ml of blasticidin S for 2 days, respectively. The PML knockout clones were screened from the resultant single colonies by immunofluorescence staining with anti-PML antibody and further analyzed by sequencing of PML cDNA from each clone to confirm faithful knockout of *Pml* (data not shown).

For establishment of YS300 deleted cell line (ΔYS), mESCs were transfected with pSpCas9(BB)-2A-Puro (PX459) V2.0 harboring three sgRNAs targeting the YS300 allele by using Lipofectamine 2000 and selected with 3 µg/ml of puromycin for 2 days. Genomic positions targeted by these sgRNAs were shown in Supplementary Fig. 5a. The resultant clones were screened by DNA-FISH with a BAC probe (B6Ng01-016L17) for YS300 (data not shown). For generation of Pml KO mESCs rescued by expression of exogenous PML gene (Pml KO + PML rescue), Pml KO cells were electroporated with both a BAC DNA (B6Ng01-109M19) encoding *Pml* locus and pCAG-H2B-iRFP carrying a puromycin resistance gene, and then selected with 2 µg/ml of puromycin for a week. The targeted clones were screened by immunofluorescence staining with anti-PML antibody (data not shown). All the targeting vectors, the sgRNA sequences, and BAC clones used in this study are listed in Supplemental table.

### Antibodies

Anti-APEX2 antibody was raised by immunization of recombinant APEX2 protein to a rabbit. Antibodies against PML (MBL, 36-1-104), PML (Santa Cruz, sc-5621), TRF1 (a gift from K. Okamoto)^46^, NANOG (eBioscience, eBioMLC-51), EZH2 (Cell Signaling Technology, AC22), DNMT3A (Imgenex, IMG-266), DNMT3B (Cell Signaling Technology, 48488S), H3K27me3 (Cell Signaling Technology, C36B11), and Digoxigenin (DIG) (Roche), α-Tubulin (Calbio, DM1A) were used for either immunostaining or western blot. Antibodies against H3K4me3 (Cosmo Bio, MABI0304), H3K27ac (Cosmo Bio, MABI0309), H3K27me3 (Cell Signaling Technology, C36B11), H3K9me3 (Diagenode, C15500003), PML (MBL, 36-1-104), HA (Sigma, H9658) and GFP (Abcam, ab290) were used for ChIP assay.

### Western blot

Proteins were resolved in SDS-PAGE gels, immunoblotted and detected as previously described^47^.

### Fluorescent staining of cells

Cells were grown on a micro-insert 4 well (Ibidi) placed on a glass-bottom dish (Matsunami) coated with iMatrix-511 (Nippi, 892021). Immunostaining was performed as previously described^48^. Appropriate secondary antibodies labeled with fluorophores such as Alexa Fluor dyes were used for secondary antibodies. Biotinylated fractions were stained with fluorescently labeled Streptavidin (StAv).

DNA-FISH was performed as previously described^49^. In brief, cells were fixed in 4% formaldehyde in phosphate buffered saline (PBS) for 10 min at 37 °C, and then incubated with PBS containing 0.5 µg/ml RNase A and 0.5% Triton X-100 for 10 min at room temperature (RT). The cells were treated with 0.1N HCl for 2 min at RT, and re-fixed with 1% formaldehyde for 5 min. After applying hybridization mixture containing 50% formamide, 10% dextran sulphate, 2x Saline Sodium Citrate buffer (SSC), 1 mg/ml mouse Cot1 DNA, 1 mg/ml polyvinyl pyrrolidone (PVP), 0.01% Triton X-100, 1 mg/ml Bovine serum albumin (BSA), and 10 ng/µl of fluorescently labeled probes to the cells, the specimens were denatured for 5 min at 80 °C, and then incubated at 37 °C overnight. After washing once with 2x SSC containing 0.01% Triton X-100 for 10 min and twice with 0.2x SSC containing 0.01% Triton X-100 for 10 min at 45 °C, the specimen was mounted in Vectashield (VECTOR, H-1200) or Prolong Diamond (Thermo Fisher, P36965). Fluorescent probes with BAC DNAs were prepared by nick translation with dATP conjugated with fluorophore or DIG using Nick Translation kit (Roche) or PCR. Probes for the ALaPed DNA and chromosome painting for the Y chromosome were prepared by PCR. Immuno-DNA-FISH was performed as previously described^48,49^. In brief, after immunostaining as described above, cells were fixed in 4% formaldehyde in PBS for 10 min at 37 °C to cross-link antibodies. DNA-FISH was subsequently performed as described above.

For Immuno-DNA-RNA-FISH analysis, RNA-FISH was first performed as previously described^48^. Briefly, cells were fixed with 4% formaldehyde in PBS for 20 min at RT, and then permeabilized with 0.5% Triton X-100 in fixative for 10 min followed by dehydration in ethanol series 70%, 100%, and air dried. The cells were incubated with hybridization mixture (50% formamide, 10% dextran sulphate, 2x SSC, 1 mg/ml PVP, 0.05% Triton X-100, 0.5 mg/ml BSA, 1 mg/ml mouse Cot1 DNA, and 10 ng/μl DIG labeled probes) at 50 °C overnight, and washed once with 2x SSC containing 0.01% Triton X-100 for 10 min and subsequently twice with 0.2x SSC containing 0.01% Triton X-100 for 10 min. Immunostaining was subsequently performed with anti-DIG and anti-PML antibodies. The cells were then fixed again with 4% formaldehyde for 10 min and processed to DNA-FISH as described above. Primers used for PCR were listed in Supplemental table. All the images were acquired by using A1R confocal laser microscope (Nikon) or Cell Voyager CV1000 confocal scanner system (Yokogawa).

### ALaP-seq

The APEX reaction was performed as previously described with some modifications^15^. 1 × 10^7^ Nanog-APEX or APEX-Pml cells were incubated with 500 µM biotin-phenol in culture medium for 10∼15 hrs at 37 °C. The cells were then treated with 1 mM H_2_O_2_ for 1 min at 37 °C to trigger biotinylation. The reaction was immediately quenched by washing the cells with pre-chilled PBS containing 10 mM sodium azide, and 10 mM sodium ascorbate. The cells were subsequently permeabilized with pre-chilled NP-40 lysis buffer (20 mM Tris-HCl pH 7.6, 1 mM EDTA, 150 mM NaCl, 0.1% IGEPAL CA-630, 10 mM sodium azide, 10 mM sodium ascorbate) on ice for 3 min. Then, fixation was performed with 1% formaldehyde in NP-40 lysis buffer for 10 min on ice and subsequently stopped by adding glycine at a final concentration of 0.125 M. The fixed cells were collected in a 15 ml tube, and washed twice with quencher solution followed by centrifugation at 1,500 rpm for 5 min. The pellet was resuspended in 500 μl of digestion buffer containing 20 mM Tris-HCl, pH 8.0, 5 mM NaCl, 2.5 mM CaCl_2_, 1x Protease Inhibitor Cocktail (Nakarai, 03969-21), and 5U of Micrococcal Nuclease (Takara, 2910A) and incubated for 10 min at 37 °C. The reaction was stopped by adding 10 mM EDTA and 5 mM EGTA. After lysis of cells by brief sonication, 500 μl of LB3 buffer (195 mM NaCl, 0.2% Na-Deoxycholate, 1% Na-Lauroylsarcosine, 5% BSA) and 100 µl of 10% Triton X-100 were added to the lysate. The lysate was centrifuged at 15,000 rpm and 4 °C for 10 min. The supernatant was mixed with 100 μl of StAv-coated magnetic beads (ThermoFisher, 65602) and incubated at 4 °C overnight with gentle rotation. After the incubation, StAv beads were washed seven times with RIPA buffer (50 mM Hepes-NaOH, pH 7.2, 500 mM LiCl, 1 mM EDTA, 0.1% IGEPAL CA-630, 0.7% Na-Deoxycholate, 1% sodium dodecyl sulfate (SDS), and 2 M Urea) and two times with TE buffer, including 0.01% Triton X-100. Chromatin was eluted by incubation in reverse crosslink buffer (50 mM Tris-HCl, pH 8.0, 10 mM EDTA, 0.1% SDS, and 50 µg/ml Proteinase K) at 65 °C for 12 hrs and purified with Monarch DNA Cleanup Columns (New England Biolabs, T1030L). For ALaP-qPCR analysis, the relative binding level of each genomic region was normalized against input. Primer sequences are listed in supplementary Table. For ALaP-seq, library was prepared using NEBNext ultra II (New England Biolabs, E7645S) and NEBNext primers (New England Biolabs, E7600S), and subsequently size selected to 150-500 bp using SPRIselect beads (Beckman, B23317). The libraries were sequenced on Illumina HiSeq3000 for 100 bp paired-end reads.

### RNA-seq

Total RNA was extracted from wild-type mESCs and Pml KOs using RNeasy Mini Kit (QIAGEN, 74104) and treated with Turbo-DNase (Ambion, AM2238). poly-A RNA was isolated from total RNA by using oligo(dT) beads (ThermoFisher, 61002) and libraries were prepared with TruSeq RNA prep kit (Illumina, RS-122-2101). Three biological replicates were prepared for each sample. Libraries were sequenced on Illumina HiSeq2500 for 100 bp single-end reads. The transcriptome data sets of PML knockdown mESCs and PML knockout MEFs were obtained from previous reports^13,45^.

### ChIP-seq

ChIP was performed as described previously^50^ with minor modifications. In brief, 2 × 10^6^ cells were fixed in 1% formaldehyde for 10 min at RT. The fixation was subsequently quenched with 0.125 M glycine. Chromatin was sonicated to 200-500 bp by using Branson Sonifier 250 (Branson Ultrasonics). The fragmented chromatin was immunoprecipitated for overnight at 4 °C using respective antibodies complexed with anti-mouse, rabbit IgG Dynabeads, or StAv beads (ThermoFisher, 11201D, 11203D and 11205D) After washing the beads, the library was prepared by ChIPmentation strategy^51^. Briefly, beads-bound chromatin was tagemented in the reaction mixture composed of 5 μl of 5x tagmentation buffer, 2.5 μl of dimethylformamide (DMF), 1.25 μl of Tn5 transposase, and 16.25 μl of nuclease-free water at 37 °C for 10 min. Chromatin was eluted by incubation in the reverse crosslink buffer at 65 °C for 12 hrs and purified with Monarch DNA Cleanup Columns. The DNA was amplified by PCR using Nextera primers^52^ and size-selected to 200-500 bp using SPRIselect beads, followed by sequencing on Illumina HiSeq3000 with 36 or 50 bp single-end reads mode. Two biological replicates were prepared for both wild-type mESCs and Pml KO.

### ATAC-seq

ATAC-seq was performed as previously described^52^. In brief, 1 × 10^5^ cells were washed with ice-cold PBS and resuspended in 50 μl of lysis buffer (10 mM Tris-HCl, pH 7.4, 10 mM NaCl, 3 mM MgCl_2_ and 0.2% IGEPAL CA-630). After centrifugation of cells at 1,500 rpm for 5 min, the pellet was resuspended in 50 μl of reaction mixture (10 μl of 5x tagmentation buffer, 5 μl of DMF, 2.5 μl of Tn5 transposase and 32.5 μl of nuclease-free water) and incubated for 30 min at 37 °C. The DNA fragments were isolated with Monarch DNA Cleanup Columns. Library amplification was performed as described for ChIP-seq. The DNA was size-selected to 200-500 bp using SPRIselect beads, followed by 100 bp paired-end sequencing on Illumina HiSeq3000. Two biological replicates were prepared for wild-type mESCs and Pml KO.

### RNA-seq analysis

For RNA-seq analysis, reads were processed with Cutadapt^53^ to trim adaptor sequences and aligned to mouse genome (UCSC mm10) using Tophat^54^. Differential expression analyses were performed using featureCounts^55^ and edgeR^56^. DEGs were selected with FDR cut off of 0.001 and |Log_2_(FC)| > 1. Gene ontology analyses for DEGs were performed using DAVID^44^.

### Treatments of deep sequencing data and peak calling for chromatin analyses

SequencereadsweretrimmedbyusingTrimGalore (https://github.com/FelixKrueger/TrimGalore) for ALaP-seq and ATAC-seq, or Trimmomatic^57^ for ChIP-seq. Trimmed reads were aligned to mouse genome (UCSC mm10) using bowtie2^58^ with parameters --very-sensitive --maxins 2000 (for paired-end). Non-uniquely mapping reads were filtered out using XS tag or a MAPQ threshold of 30. PCR duplicates and ENCODE blacklisted regions^59^ were removed using Picard Tools (http://broadinstitute.github.io/picard/) and bedtools^60^.

ALaP-seq peaks were independently identified for two biological replicates using Model-based Analysis for ChIP Sequencing v2.0 (MACS2)^61^ with FDR cut off of 0.001, and the overlapped peaks between the replicates were used for downstream analyses. ChIP-seq peaks for SUMO-1 and SUMO-2/3 were obtained from GSE99009^28^, and the coordinates were converted from mm9 to mm10 using liftOver function from the rtracklayer package^62^. The normalized read coverages for ChIP-seq and ATAC-seq were computed by using deeptools bamCoverage with option --normalizeUsing RPKM, and visualized with Integrative Genomic Viewer^63,64^. The read Coverage for ALaP-seq was calculated by using bamCompare with option --scaleFactorsMethod SES --operation subtract. To compensate the coverage between sex chromosomes and autosomes, the coverage scores on Chr X and Y were scaled twice.

### Peak annotation

Peaks of ALaP-seq were annotated for the genomic elements and associated genes using Homer^65^. Enhancer regions in mouse ES cells were obtained from a previous study^66^, in which the enhancers were predicted based on ChIP-seq data of enhancer signatures, including p300, MED12, NIPBL, and H3K4me1. Its coordinate was lifted over to mm10.

### Enrichment analyses of sequence reads

Metagene analyses for enrichment around TSSs, ALaP peaks, CpG islands were performed by using EaSeq^67^ and Agplus^68^. Pairwise Pearson’s correlations between ALaP-seq, ChIP-seq, and ATAC-seq within TSS+/-500 bp were computed and plotted as heatmaps using deeptools multiBigwigSummary and plotCorrelation^64^. Whole genome bisulfite sequencing data in mouse ES cell (E14) cultured in 2i serum/LIF condition was downloaded from GSM1027572^69^ and its coordinate was lifted over to mm10.

### Analyses of ALaP enrichment within 300 kb genomic block

For enrichment analyses of ALaP-seq in 300 kb genomic block, we analyzed ALaP-seq data to mimic DNA-FISH features, including masking of repetitive sequences with Cot1 DNA. Reads were aligned to the repeat masked mouse genome (UCSC mm10) using bowtie2 with parameter -N1. The blacklisted regions^59^ and gray-listed regions prepared by chipseq-greylist (https://bioconductor.org/packages/release/bioc/html/GreyListChIP.html), both of which are likely to give false positives, were removed out. Filtering out of reads displaying multiple alignment was omitted here. Hence, the multiple-mapped reads are randomly aligned only once to only one of the selected alignment positions. Number of reads in each 300 kb genomic block throughout mouse genome was counted by using featureCounts, and then analyzed by using edgeR. Computed Log_2_ fold change for each block compared with H_2_O_2_-omitted condition, a negative control, was plotted along with each chromosome using Sushi^70^.

### Comparison between ALaP-seq and DNA-FISH with BAC DNAs

To analyze correlation between ALaP-seq enrichments and mean 3D distances (µm) from the closest PML body to DNA-FISH spot, number of reads aligned to genomic region encoded by each BAC clone was counted using featureCounts, and then normalized by total mapped reads. The ALaP enrichment score for each BAC region was calculated by dividing the normalized read counts by that of the experimental negative control where H_2_O_2_-treatment was omitted. 3D distance between gravity center of PML bodies and DNA-FISH foci was calculated as described below.

### Dot plot analysis of YS300

Dot plot analysis for similarity of genomic sequence within YS300 (chrY;110001-363557) was performed by using YASS^71^ with default parameters.

### Quantitative real-time PCR

For gene expression analysis, total RNA was isolated using a NucleoSpin RNA kit (Takara, 740955). Reverse transcription was carried out using PrimeScript II (Takara, 6210A) and random hexamers. Real-time PCR was performed with Thunderbird SYBR qPCR Mix (Toyobo, QPS-201) or KOD SYBR qPCR Mix (Toyobo, QKD-201) on a LightCycler 480 Real-Time PCR System (Roche). The relative expression level of each transcript was normalized by *Gapdh* expression levels. Primer sequences are listed in Supplementary table. For ChIP and ALaP qPCR analyses, telomeric sequences and the other genomic regions were quantified using PowerUp SYBR Green Master Mix (ThermoFisher, A25742) and Thunderbird SYBR qPCR Mix, respectively.

### Bisulfite sequencing

Genomic DNA was extracted from wild-type, Pml KO, or ΔYS mESCs, and subjected to sodium bisulfite conversion using EZ DNA Methylation-Gold kit (Zymo Research, D5005). Promoter regions of selected genes were amplified by PCR using KOD-Multi&Epi (TOYOBO, KME-101) and cloned into TA cloning vector pTAC-2 (Biodynamics Laboratory). The inserts were sequenced and analysed using Quantification Tool for Methylation Analysis^72^.

### Quantification of 3D distances between PML bodies and DNA-FISH foci

Image processing and data analyses were performed on the in-house developed applications in C, Ruby and R. Prior to all the processes, image resolutions along Z-axis of image stacks were first interpolated by Lanczos-5 method in accordance with that of X- and Y-axes. The shape of the nuclei was segmented by applying 3D watershed algorithm to the fluorescence of gelGreen or DAPI. The signal of PML bodies and chromatin regions within the individual nucleus were segmented by threshold values determined by Renyi’s entropy method^73^, and their positional information was organized as SQLite3 database. Euclidean distance for every pair of a PML body and gene locus within the nuclei were calculated based on the gravity center of objects. The shortest distance was determined for each FISH focus based on the rank of a pair which showed the most adjacent spatial relationship.

### Relative quantification of size of PML bodies adjacent to genetic loci

The volume was calculated for each PML body. To compensate different number of PML bodies in each nucleus, the ranks of the volume were normalized by number of PML bodies, providing the quantile ranking of the volume in each nucleus.

### DNA-FISH analysis in ΔYS mESCs

To quantify the intensity of DNA-FISH for YS300 and the *Uty* locus in wild-type and ΔYS mESCs, all the images were binarized with appropriate thresholds. Total intensity of each fluorescence spot was quantified using ImageJ. The fluorescent intensity of DNA-FISH was normalized by median of corresponding score in wild-type mESCs. The center-center 3D distance between PML bodies and YS300 or the *Uty* was calculated using ImageJ plugin DiAna^74^. 3D colocalization between PML bodies and TRF1, or DNA-FISH foci was also analyzed with DiAna.

### Averaged image and radial distribution analysis

Cells were co-stained for PML and either DNMT3A, DNMT3B, or EZH2. Averaged image projections were prepared as described previously^75^. In brief, PML bodies were identified in individual z-stacks through appropriate intensity thresholds, and centered along a box of size (l = 2.76 μm). Images for all channels at the l × l square centered at each PML body at every corresponding z-slice were cropped and combined to quantify average fluorescent intensity, providing averaged projection images centered at PML bodies.

### Statistical analysis

Statistical comparisons for Fig. 1d was performed using Dunnett test. For Fig. 3b, c, Fig. 5g, i and Supplementary Fig. 8d, *p* values were calculated using Mann–Whitney U test. Significances for Fig. 5h and Supplementary Fig. 1d were determined using two-sided t test with unequal variance and one-way ANOVA with Tukey HSD test, respectively.

## Acknowledgements

We appreciate K.Okamoto for sharing an anti-TRF1 antibody, M.Okano for providing the Dnmt^TKO^ mESCs, Feng Zhang for a gift of Addgene plasmid no. 62988, P.Kalitsis for a plasmid pYmin2.3a. We also appreciate K.Yamaguchi, A.Akita, M.Matsumoto for their technical supports for RNA-seq, K.Naruse and A.Kato for Sanger sequencing, NYU Center for Genomics and Systems Biology GenCore and NIBB Collaborative Research Program for their sequencing services, NIBB Data Integration and Analysis Facility for the computational resources, and NIBB Bioimaging Facility for their imaging supports. M.E.Torres-Padilla and A.Fadloun for critical reading of the manuscript. This work was supported by JSPS KAKENHI 16H06279 (PAGS) (Y.M.), 15H05586 (Y.M.), 18K14632 (M.K.), 18K06272 (K.K.), Okazaki Orion Project (Y.M.), Tomizawa Fund (Y.M.), Astellas Foundation (Y.M.), JST/PRESTO (Y.M.), and Nakajima Foundation (Y.M.). K.M. is a recipient of JPSP postdoctoral fellowship (2018-2019).

## Author contributions

K.M. and Y.M conceived and designed the study and wrote the manuscript, with the assistance and final approval of all authors. K.M. performed most of the experiments and analyzed the data. K.K. performed imaging analyses. C.S performed plasmid DNA constructions and establishment of mESC lines. T.F. synthesized biothin-phenol. Y.O. and S.S. contributed to the initial setup for ALaP-seq and RNA-seq.

